# Genetic disruption of the Blood Brain Barrier leads to protective barrier formation at the Glia Limitans

**DOI:** 10.1101/2020.03.13.990762

**Authors:** Pierre-Louis Hollier, Sarah Guimbal, Pierre Mora, Aïssata Diop, Lauriane Cornuault, Thierry Couffinhal, Alain-Pierre Gadeau, Marie-Ange Renault, Candice Chapouly

## Abstract

Recent work demonstrated that Central Nervous System (CNS) inflammation induces endothelial Blood Brain Barrier (BBB) opening as well as the formation of a tight junction barrier between reactive astrocytes at the Glia Limitans. We hypothesized that these two barriers may be reciprocally regulated by each other state and further, that the CNS parenchyma may acquire protection from the reactive astrocytic Glia Limitans not only in neuro-inflammation but also when BBB integrity is compromised under resting condition, without pathology. Previous studies identified Sonic hedgehog (Shh) astrocytic secretion as implicated in stabilizing the BBB during neuropathology and we recently demonstrated that desert hedgehog (Dhh) is expressed at the BBB in adults.

Here we unraveled the role of the morphogen Dhh in maintaining BBB tightness and, using endothelial Dhh knockdown as a model of permeable BBB, we demonstrated that a double barrier system comprising both the BBB and Glia Limitans, is implemented in the CNS and regulated by a crosstalk going from endothelial cell to astrocytes.

First, we showed that, under neuro-inflammatory conditions, Dhh expression is severely down regulated at the BBB and that Dhh is necessary for endothelial intercellular junction integrity as Dhh knockdown leads to CNS vascular leakage. We then demonstrated that, in Dhh endothelial knockout (Dhh^ECKO^) mice which display an open BBB, astrocytes are reactive and express the tight junction Claudin 4 (Cldn4) and showed that astrocytes can respond to signals secreted by the permeable endothelial BBB by becoming reactive and expressing Cldn4. To examine the consequences of the above results on disease severity, we finally induced multiple sclerosis in Dhh^ECKO^ mice versus control littermates and showed that the pathology is less severe in the knockout animals due to Glia Limitans tightening, in response to BBB leakage, which drives inflammatory infiltrate entrapment into the perivascular space. Altogether these results suggest that genetic disruption of the BBB generates endothelial signals capable of driving the implementation of a secondary barrier at the Glia Limitans to protect the parenchyma. The concept of a reciprocally regulated CNS double barrier system has implications for treatment strategies in both the acute and chronic phases of multiple sclerosis pathophysiology.

## Introduction

In a healthy individual, the CNS parenchyma is protected from the peripheral circulation by the blood-brain barrier (BBB) which tightly regulates the entry and exit of soluble factors and immune cells^1^. Importantly, during multiple sclerosis, the abnormal permeability of the BBB allows penetration into the CNS parenchyma of inflammatory cells and soluble factors such as autoantibodies, cytokines and toxic plasma proteins which drive lesion formation and acute disease exacerbation^2,3^. Therefore identifying key mechanisms that promote BBB tightness is currently considered as a main strategy to control leukocyte and humoral entry, preventing disease progression and severity in multiple sclerosis.

Previous studies identified the Hedgehog (Hh) pathway as a new regulator of vessel integrity in multiple sclerosis, HIV and stroke, showing its implication in stabilizing the BBB^4–7^ and we recently demonstrated that Dhh is expressed at the BBB in adults^8^. Dhh, together with Shh (sonic hedgehog) and Ihh (indian hedgehog), belongs to the Hh family identified nearly 4 decades ago in drosophila as crucial regulators of cell fate determination during embryogenesis^9^. The interaction of Hh proteins with their specific receptor patched-1 (Ptch1) derepresses the transmembrane protein smoothened (Smo), which activates downstream pathways, including the Hh canonical pathway leading to the activation of gli family zinc finger (Gli) transcription factors and the so-called Hh non canonical pathways, which are independent of Smo and/or Gli^10^.

Interestingly, recent discovery showed that considering the BBB only as a vascular structure may be a simplistic and inaccurate vision. Indeed, a substantial intercellular communication occurs between vascular cells and the adjoining glia^11^. It is more proper to consider CNS barrier structure as a multicellular neurovascular unit that includes pericytes^12^, astrocytes^13^ as well as the blood vessels themselves. More specifically, to enter the CNS from the vasculature, soluble factors and immune cells must initially traverse the BBB. Then, they circulate within the perivascular space (PVS), a region surrounding the basal surface of the endothelial cell wall to reach the CNS parenchyma by going through a network of astrocytes termed the Glia Limitans^14,15^. While it is now well established that BBB breakdown leads to soluble factor and inflammatory cell infiltration into the PVS during neuropathology^4^, the role of the Glia Limitans appears trickier. Indeed, astrocytes, described as reactive, are emerging as “Dr Jekyll and Mister Hyde” cells, having complex roles in both recruiting and restricting neuro-inflammatory infiltration^16^. Reactive astrocyte behavior is determined in a context-specific manner by signaling events that vary with the nature and severity of CNS insults. Specifically, in multiple sclerosis as well as Alzheimer’s and Parkinson’s diseases, it has been shown that reactive astrocytes, on one hand, produce pro-inflammatory and pro-permeability factors^17–19^ and on the other hand, neuro-protective factors^2^. Strikingly, the results we published in 2017 raised considerable attention on a new property of reactive astrocytes: the expression of tight junction proteins (notably Claudin4 (Cldn4)) under inflammatory conditions^20^.

The first objective of our study is to decipher the role of the morphogen desert hedgehog (Dhh) in maintaining BBB tightness. The second objective is to demonstrate that a double barrier system comprising both the BBB and Glia Limitans, is implemented in the CNS and regulated by a crosstalk going from endothelial cell to astrocytes, using endothelial Dhh knockdown as a model of permeable BBB.

In this study, we first demonstrated that endothelial Dhh expression is down regulated during neuro-inflammation and necessary to maintain BBB tightness. We then showed that BBB opening, induced by Dhh knockdown, drives astrocyte Cldn4 expression, conferring barrier properties to the Glia Limitans which result in the PVS entrapment of plasmatic protein and inflammatory cell both physiologically and during pathology. Altogether these data identify the gliovascular unit as a double barrier system whom function is controlled by the crosstalk between endothelial cell and astrocyte.

In conclusion, this work strengthen the concept of CNS double barrier system, unveiling how signals at the endothelium drive astrocyte barrier properties to protect the parenchyma during neuropathology. Consequently, taking into account both components of the gliovascular unit is of translational interest and could open the way for new therapeutic strategies notably to limit progressive multiple sclerosis pathology.

## Results

### Desert hedgehog (but not Sonic and Indian hedgehog) is expressed by CNS microvascular endothelial cells and downregulated during chronic neuro-inflammation

First, we showed that Dhh is expressed at the BBB both *in vitro* using mouse CNS MECs* (Figure 1, A) and *in vivo* using human cortical sections from healthy donors (Figure 1, B). Shh and Ihh are not expressed in the healthy CNS confirming the single cell sequencing data from the Betsholtz’s lab^25^ and Dhh is known to be stored intracellularly as well as being secreted^7^. Therefore, we here infer the CNS endothelial cells as the source of Dhh within the gliovascular unit. Next, we demonstrated that Dhh is severely down regulated at the BBB under inflammatory conditions both *in vitro* using HBMECs treated with IL-1β (one of the main pro-inflammatory cytokine implicated in multiple sclerosis pathophysiology) (Figure 1, C) and *in vivo* (Figure 1, F) using a pre-clinical model of multiple sclerosis (MOG_35-55_) to induce chronic neuro-inflammation in C57BL/6 mice. Dhh down regulation at the BBB is associated to the up regulation of endothelial activation markers Icam1 (Figure 1, D, G) and Vcam1 (Figure 1, E, H) and to the down regulation of tight junctions (Cldn5 and Zo1) (Figure 1, I-J).

**Figure 1:**
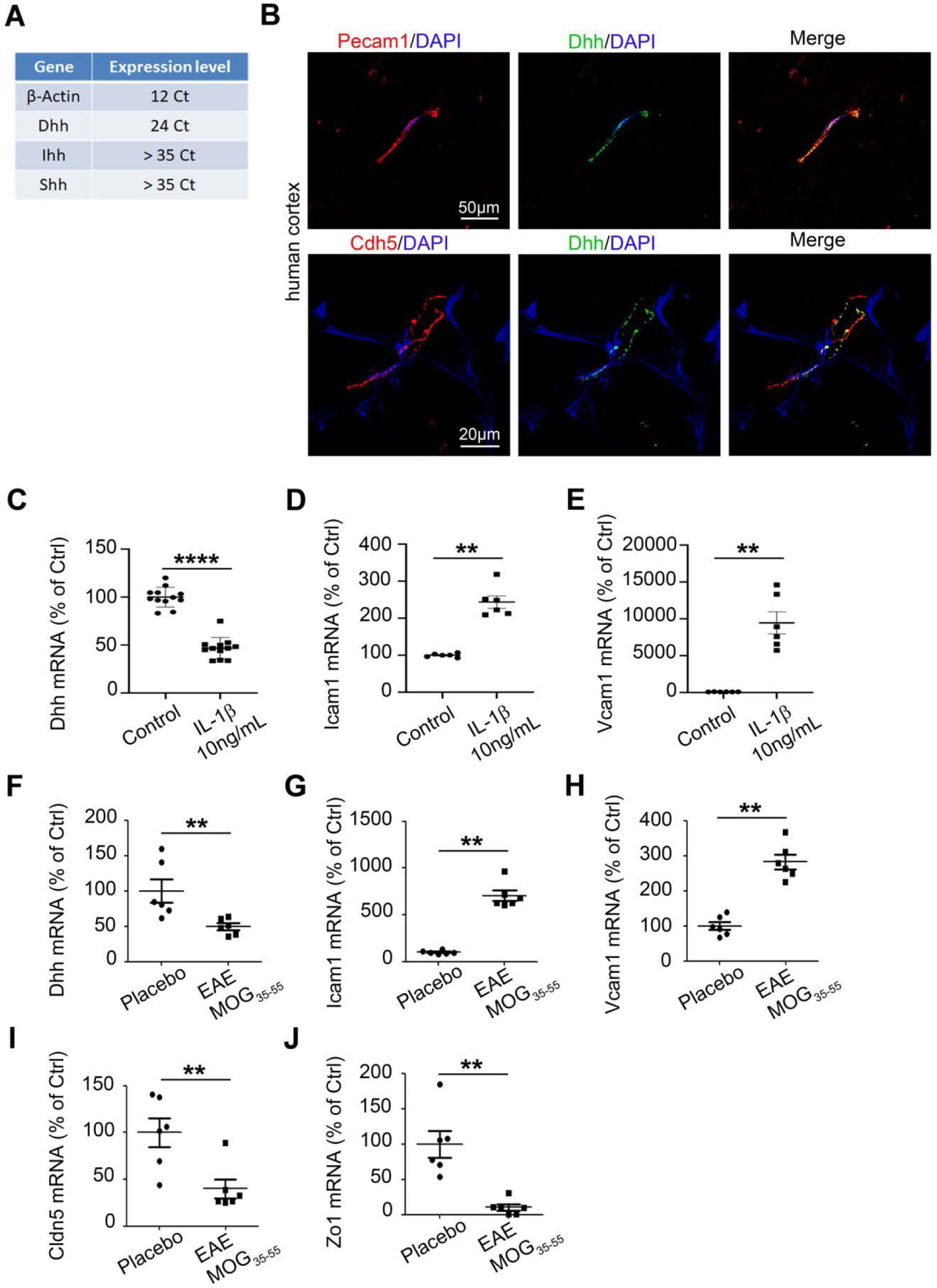
Desert Hedgehog is expressed by CNS microvascular endothelial cells and downregulated during chronic neuro-inflammation: **(A)** Primary Brain Microvascular Endothelial Cells (BMECs) were isolated from 12 weeks old C57Bl/6 mice and Dhh, Ihh and Shh expression were quantified by RT-qPCR (cycle threshold mean values).β-Actin is used as a reference. **(B)** Human cortical sections from healthy donors were obtain from the NeuroCEB biobank and immunostained with anti-Cdh5 (in red), anti-Pecam1(in red) and anti-Dhh (in green) antibodies. Nuclei were stained with DAPI (in blue). **(C-E)** HBMECs were cultured until confluency and starved for 24h. HBMECs were then treated with PBS (control condition) or IL1β 10 ng/mL for 24h and **(C)** Dhh, **(D)** Icam1 and **(E)** Vcam1 expression were quantified by RT-qPCR. **(F-H)** 12 weeks old C57Bl/6 females (6 animals per group) were induced with MOG_35-55_ EAE versus Placebo. At day 16 post induction, mice were sacrificed and Spinal Cord Microvascular Endothelial Cells were isolated. **(F)** Dhh, **(G)** Icam1, **(H)** Vcam1 **(I)** Cldn5 and **(J)** Zo1 expression were measured via RT-qPCR in both groups (MOG_35-55_ versus placebo).

Altogether these data identify Dhh as the only Hh expressed in adults at the endothelial BBB. Moreover they highlight the fact that Dhh expression is down regulated at the BBB during neuro-inflammatory pathology.

*\* It is important to note that CNS endothelial cells are from a pooled source including both brain and spinal cord tissues. Therefore the resulting cell cultures/lysates may be heterogeneous in their use of Dhh. This remark applies to Figure 1, Figure 2, Figure 3 and Figure 4.*

**Figure 2:**
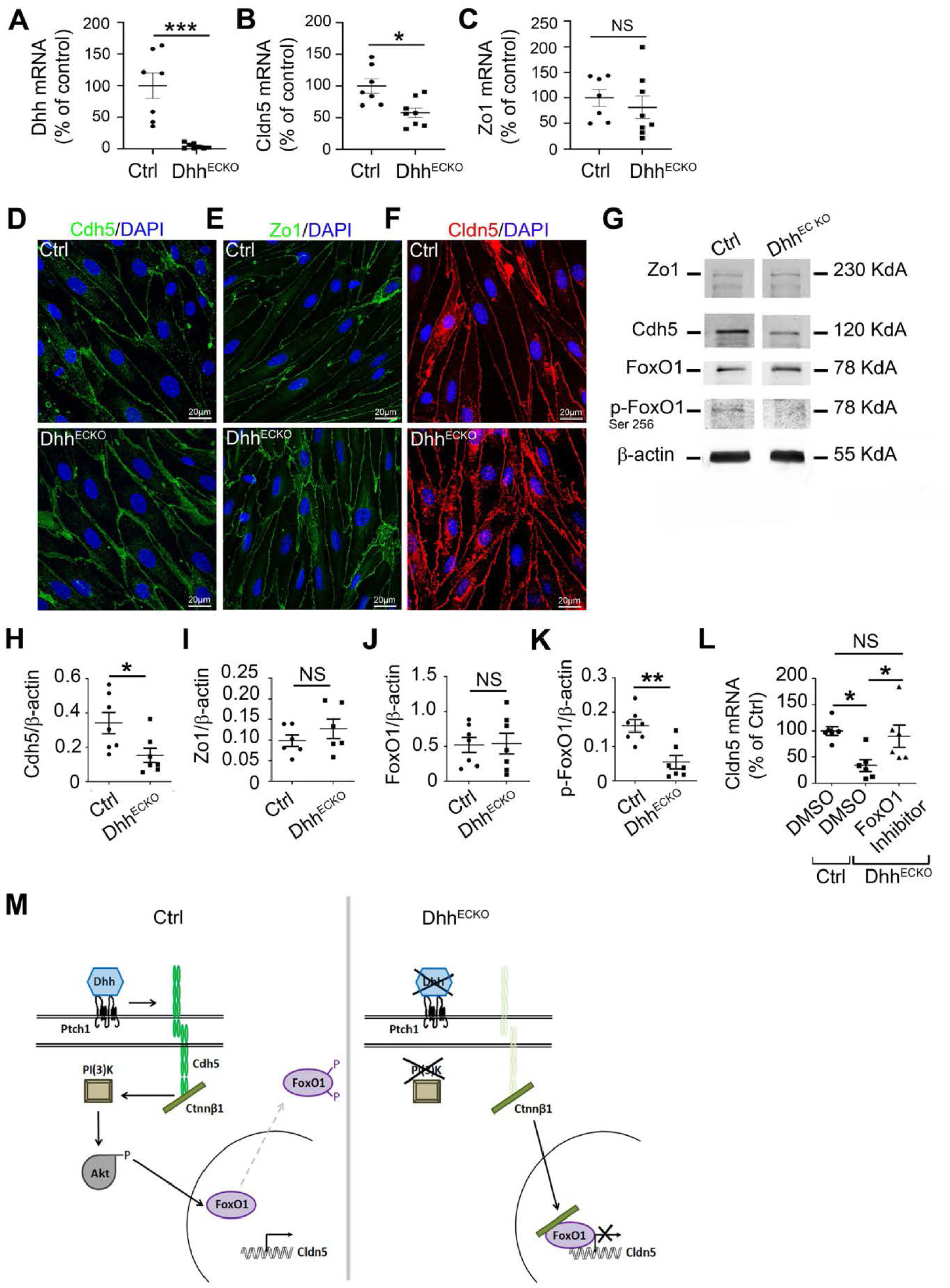
Endothelial-specific Dhh inactivation induces adherens and tight junction downregulation through stimulation of FoxO1 transcriptional activity: **(A-C)** CNS MECs were isolated from Dhh^ECKO^ and control mice and **(A)** Dhh, **(B)** Cldn5 and **(C)** Zo1 expression were quantified by RT-qPCR. **(D-F)** primary BMECs from Dhh^ECKO^ and control mice were isolated and cultured on Lab-Tek®. **(D)** Cdh5 (in green), **(E)** Zo1 (in green) and **(F)** Cldn5 (in red) localizations were evaluated by immunofluorescent staining of a confluent cell monolayer. Nuclei were stained with DAPI (in blue). The experiment was repeated 3×. **(G-L)** CNS MECs were isolated from Dhh^ECKO^ and control mice and **(G, H)** Cdh5, **(G, I)** Zo1, **(G, J)** FoxO1, **(G, K)** and p-FoxO1 expression were quantified by western blot. **(L)** BMECs were isolated from Dhh^ECKO^ mice and control mice, cultured until confluency and starved for 24h. Control BMECs were then treated with DMSO and Dhh^ECKO^ BMECs with DMSO versus an inhibitor of FoxO1 (AS1842856). **(L)** Cldn5 expression was then quantified by RT-qPCR. **(M)** Summary outline of the up regulation of endothelial junctions by the morphogen Dhh through the inhibition of FoxO1 transcriptional activity. \**P*≤0.05, *** *P*≤0.001 Mann-Whitney U test. \**P*≤0.05 Kruskal Wallis test

**Figure 3:**
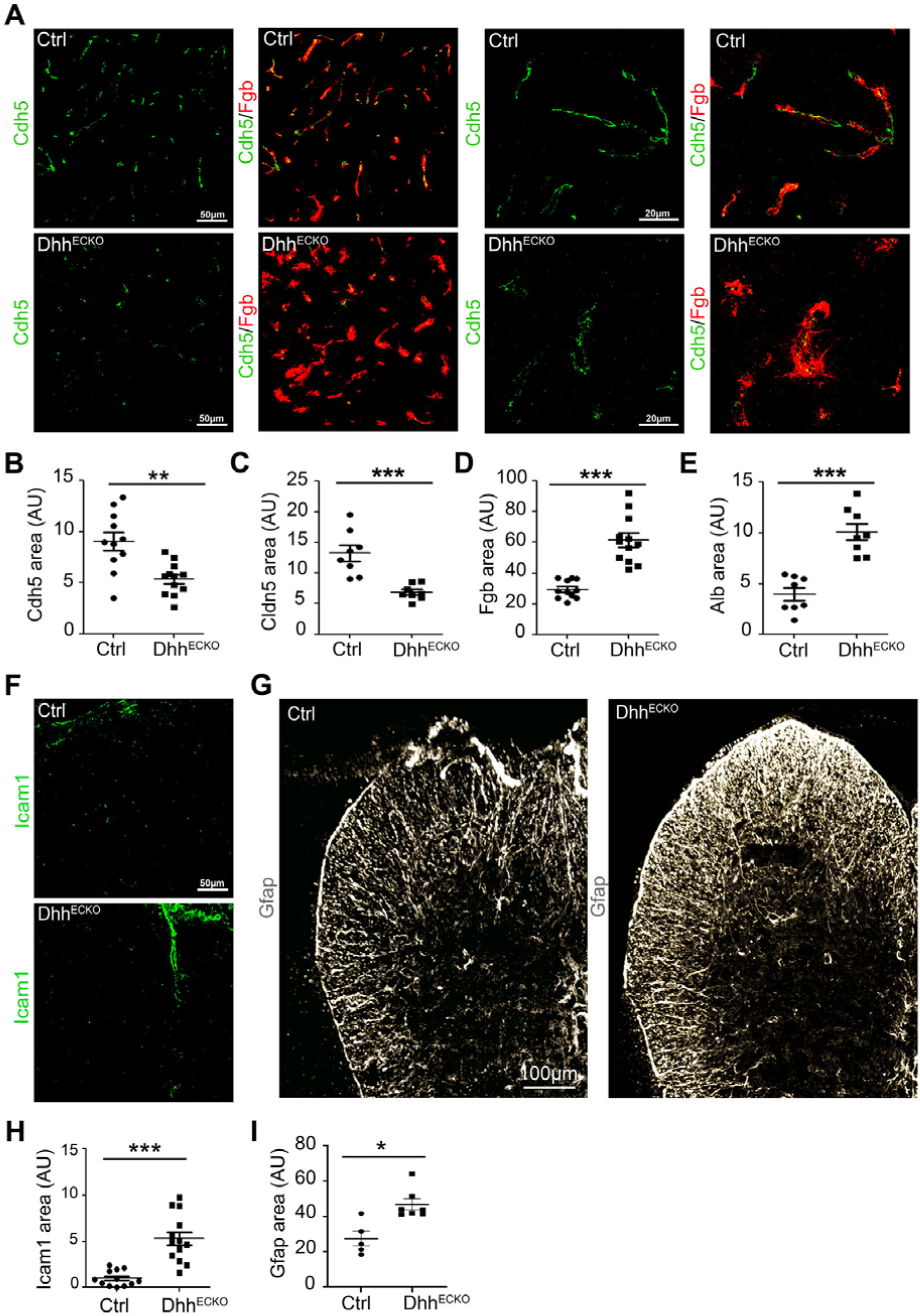
Endothelial-specific Dhh inactivation induces BBB permeability associated to endothelial and astrocytic activition in vivo: **(A-E)** Spinal cord sections were harvested from Dhh^ECKO^ mice and littermate controls and immunostained with anti-Cdh5, anti-Cldn5, anti-Fgb and anti-Alb antibodies. **(A)** Representative Cdh5 and Fgb staining were shown. **(B)** Cdh5 **(C)** Cldn5 **(D)** Fgb and **(E)** Alb positive areas were quantified (Dhh^ECKO^ *n* = 11, WT *n* = 11). **(F-I)** Spinal cord sections were harvested from Dhh^ECKO^ mice and littermate controls and immunostained with anti-Icam1 (in green) and anti-Gfap (in grey) antibodies. **(F-G)** representative Icam1 and Gfap staining were shown. **(H)** Icam1 and **(I)** Gfap positive areas were quantified. \**P*≤0.05, \*\**P*≤0.01, *** *P*≤0.001 Mann-Whitney U test.

**Figure 4:**
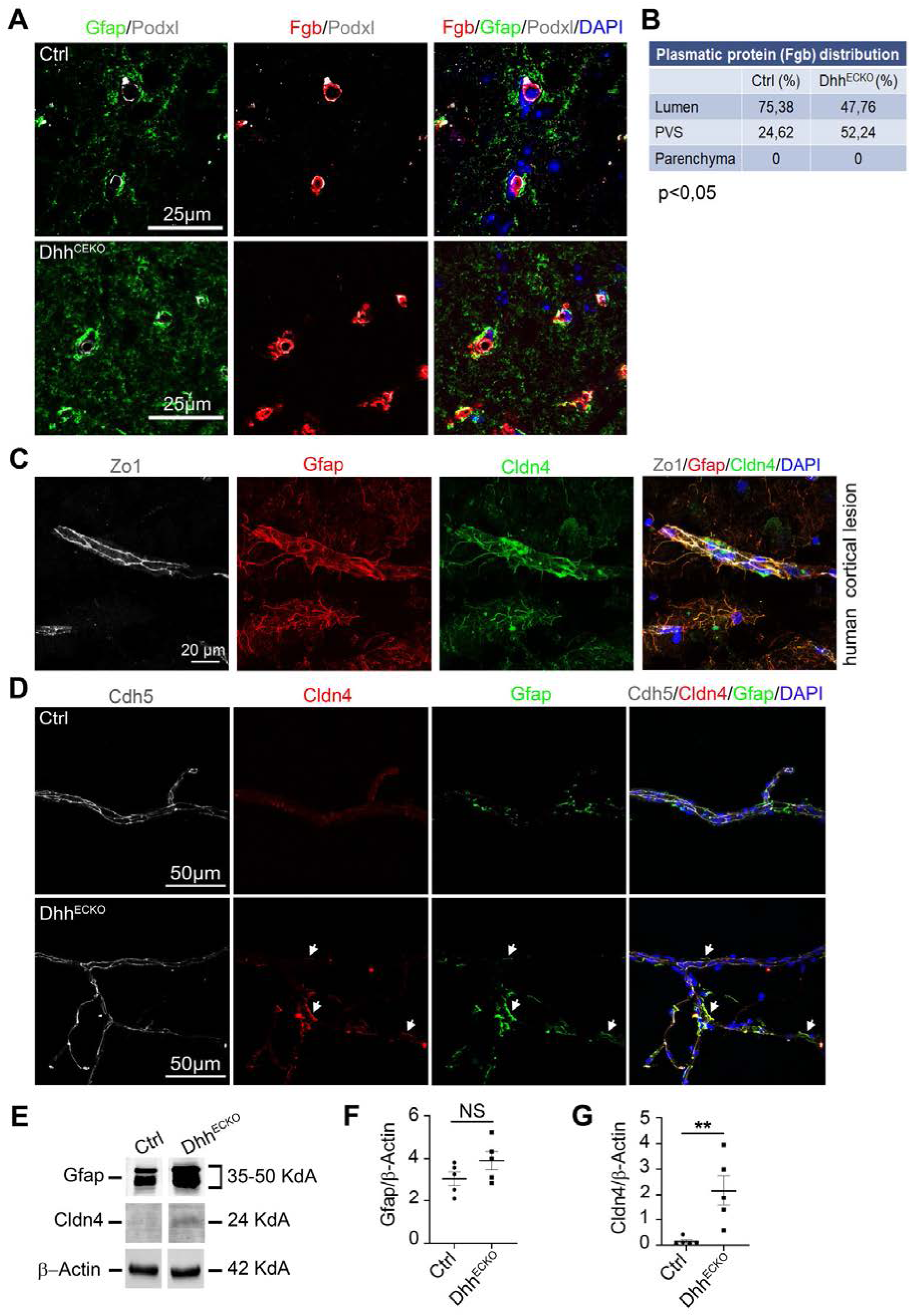
BBB breakdown is sufficient to induce a secondary CNS protective barrier at the Glia Limitans. **(A-B)** Spinal cord sections were harvested from Dhh^ECKO^ mice and littermate controls and **(A)** immunostained with anti-podxl (in grey), anti-gfap (in green) and anti-Fgb (in red) antibodies (nuclei were stained with DAPI (in blue)) and **(B)** the distribution of Fgb within the lumen, PVS and parenchyma, was quantified. We quantified the amount of Fgb in the PVS and parenchyma and inferred from it the amount of Fgb contained in the lumen. **(C)** Human active multiple sclerosis cortical lesions were obtain from the NeuroCEB biobank and immunostained with anti-Zo1 (in grey), anti-Gfap (in red) and anti-Cldn4 (in green) antibodies. Nuclei were stained with DAPI (in blue). **(D-G)** Neurovascular units were isolated from Dhh^ECKO^ and control mouse CNS and **(D)** immunostained with anti-Cdh5 (in grey), anti-Cldn4 (in red) and anti-Gfap (in green) antibodies (nuclei were stained with DAPI (in blue)). **(E-G)** Gfap and Cldn4 expression level were quantified by western blot on Dhh^ECKO^ and control mouse neurovascular unit lysates. (Dhh^ECKO^ *n* = 5, WT *n* = 5). \**P*≤0.05, \*\**P*≤0.01, Mann-Whitney U test.

### Endothelial-specific Dhh inactivation induces adherens junction Cdh5 and tight junction Cldn5 down regulation ex vivo

To test the importance of endothelial Dhh expression at the BBB, we conditionally disrupted Dhh expression in endothelial cells, and examined the consequences on BBB integrity.

We first verified the efficiency of the knockout by measuring Dhh expression in primary CNS MECs isolated from Dhh^ECKO^ and littermate controls and showed that Dhh expression is strongly down regulated in the knockout mice (Figure 2, A). Moreover we showed that Cdh5, Cldn5 and Zo1 junctions are disorganized in Dhh^ECKO^ endothelial cells: Cdh5 junctions are duplicated and blurry and Cldn5 and ZO1 junctions present a Z organisation (Figure 2, D-F). Cdh5 and Cldn5 but not Zo1 are down regulated in Dhh^ECKO^ CNS MECs compared to controls (Figure 2, B-C, G-I).

We concluded that endothelial Dhh expression is necessary to maintain endothelial adherens and tight junction expression at the BBB.

### Dhh induces Cldn5 up regulation through the inhibition of FoxO1 transcriptional activity in vitro

We previously identified Dhh as a factor facilitating the interaction between Cdh5 and Ctnnβ1 in endothelial cells^8^ and a paper published by Dejana’s lab in 2008 showed that Cdh5 interacts with Ctnnβ1 to inhibit transcription factor FoxO1^26^ via a PI(3)K–Akt dependent phosphorylation, which thereby upregulate the expression of the endothelial tight junction protein Cldn5^26^. Therefore, we next measured the expression level of the phosphorylated form of FoxO1 (p-FoxO1) in CNS MECs from Dhh^ECKO^ and control mice and demonstrated that p-FoxO1 is down regulated in Dhh^ECKO^ mice compared to controls (Figure 2, G, J-K). We then treated CNS MECs from Dhh^ECKO^ mice with a cell permeable inhibitor of the transcription factor FoxO1 (AS1842856) which blocks the transcription activity of FoxO1 and measured the expression level of Cldn5. We demonstrated that Cldn5 expression, in Dhh^ECKO^ CNS MECs treated with the inhibitor of FoxO1, returns to its expression level in control CNS MECs, unlike the Dhh^ECKO^ CNS MECs treated with DMSO (Figure 2, L).

We concluded that endothelial autocrine Dhh expression at the BBB maintains the pool of Cdh5-Ctnnβ1 signaling in endothelial cells which promotes Cldn5 expression through the inhibitory phosphorylation of the transcription factor FoxO1 (Figure 3, M).

### Endothelial-specific Dhh inactivation induces BBB permeability associated to endothelial and astrocytic activation in vivo

*In vivo*, on spinal cord sections from control and Dhh^ECKO^ mice, we confirmed that the adherens junction Cdh5 and tight junction Cldn5 are down regulated under resting conditions (Figure 3, A-C) and demonstrated that it is associated to an increase accumulation of serum proteins (Fgb and Alb)^27^ around the vessels suggesting BBB opening (Figure 3, A, D-E).

We then analyzed the activation status of both the BBB and Glia Limitans in Dhh^ECKO^ and control littermates. Using spinal cord sections, we revealed that Icam1, a marker of endothelial activation, is up regulated in Dhh^ECKO^ mice compared to littermate controls (Figure 3, F, H) and associated to the up regulation of Gfap, a marker of astrocyte activation (Figure 3, G, I).

Here, we highlighted that Dhh expression at the endothelium controls BBB tightness and demonstrated that endothelial-specific Dhh inactivation at the BBB drives endothelial and astrocyte activation.

### BBB breakdown is sufficient to induce a secondary CNS protective barrier at the Glia Limitans

As we already demonstrated in Figure 3, Dhh^ECKO^ mice display BBB leakage whereas control littermates feature a tight BBB.

Although we demonstrated (Figure 3, A, D-G) BBB leakage in Dhh^ECKO^ mice, we noticed that infiltrating plasmatic proteins are concentrated around the vascular area and not seamlessly distributed within the parenchyma. To verify this observation, we quantified the distribution of Fgb in the 3 compartments (lumen, PVS and parenchyma). Specifically, in control mice, there is no significant endothelial permeability with more than 75% of Fgb contained in the lumen of blood vessels, 25% segregated in the PVS area limited by the astrocytic endfeet (Gfap antigen) on one side and the vessel wall (Podxl antigen) on the other side, and none found in the parenchyma (Figure 4, A-B). In Dhh^ECKO^ mice, vascular leakage is significant but what is striking is that 52 % of Fgb is contained into the PVS while none is found in the parenchyma, highlighting the presence of a secondary barrier at the Glia Limitans. (Figure 4, A-B).

We previously published that reactive astrocytes express tight junctions, notably Cldn4, under inflammatory conditions, in a mouse model of multiple sclerosis (EAE)^20^. Here, we highlighted that this data is relevant to Human since Cldn4 is also expressed at the Glia Limitans in active cortical lesions from multiple sclerosis patients (Figure 4, C). Based on the above results, we investigated if the PVS entrapment of plasmatic proteins observed in Dhh^ECKO^ mice is linked to the expression of the tight junction Cldn4 at the Glia Limitans in response to BBB permeability (Figure 4, D-G). We showed that Cldn4 is expressed at the Glia Limitans in Dhh^ECKO^ mice but not control littermates (Figure 4, D, E, G) using isolated gliovascular unit samples.

Altogether these results suggest that, in Dhh^ECKO^ mice, spontaneous BBB permeability leads to the establishment of a physical barrier at the Glia Limitans characterized by the expression of the tight junction Cldn4. Therefore, in Dhh^ECKO^ mice, astrocytic endfeet at the Glia Limitans, are « preconditioned” to form a secondary barrier protecting the parenchyma.

### Endothelial signals can drive astrocyte barrier properties at the Glia Limitans

Given the above results, we wanted to determine if astrocyte barrier formation requires signals from the endothelial BBB or from the plasmatic protein perivascular infiltrate. To do so, we first study *in vitro* the response of human astrocytes (NHA) to HBMECs conditioned media *versus* plasmatic proteins from healthy donors. HBMEC used to produce the conditioned media were treated with either the osmotic agent Mannitol or the pro-permeability factor VegfA to induce BBB breakdown through various methods.

We demonstrated that Gfap (Figure 5, A, E-I) Aldh1l1 (Figure 5, B) (markers of astrocyte reactivity) and Cldn4 expression (Figure 5, D, A-H, J) are up regulated, in the NHA treated with HBMEC conditioned media but not in the NHA treated with plasma from healthy donors. Vim (marker of astrocyte reactivity) expression level was not modulated in any condition (Figure 5, C).

**Figure 5:**
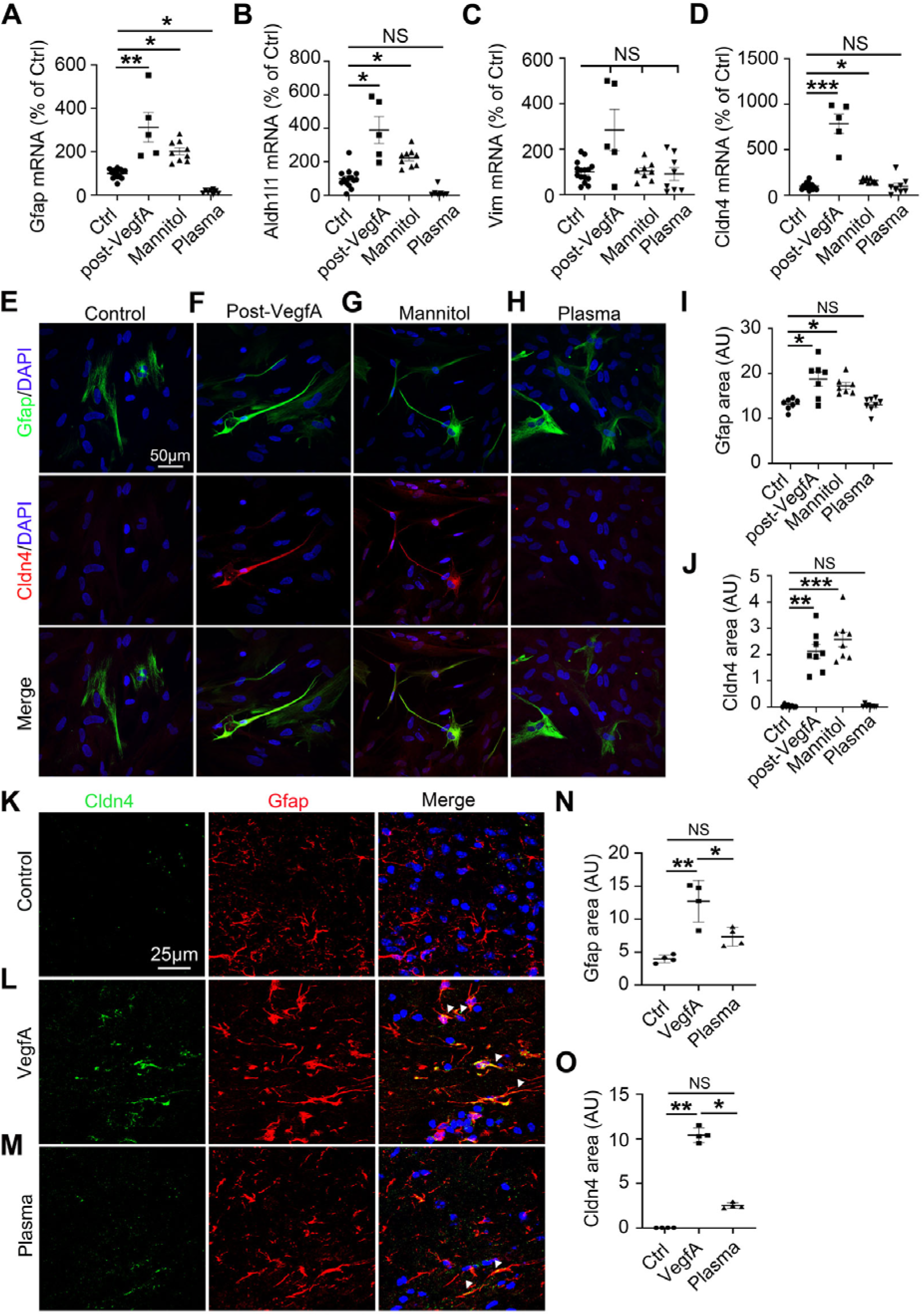
Endothelial signals can drive astrocyte barrier properties at the Glia Limitans: **(A-K)** NHA were cultured until confluency and starved for 24h. NHA were then treated for 24h with HBMEC medium from untreated cells (control condition), conditioned media form HBMECs treated with VegfA, conditioned media form HBMECs treated with Mannitol or HBMEC medium with 20% plasma from healthy donors (Mannitol and VegfA were washed out of the HBMEC cultures before the medium was used to treat the NHA cultures). **(A)** Gfap, **(B)** Aldh1l1, **(C)** Vim and **(D)** Cldn4 expression were quantified by RT-qPCR. **(E-H)** Gfap (in green) and **(E)** Cldn4 (in red) localizations were evaluated by immune-fluorescent staining of a confluent NHA monolayer. Nuclei were stained with DAPI (in blue). The experiment was repeated 3×. **(I-J)** Gfap and Cldn4 positive areas were then quantified. **(K-O)** Cerebral cortices of 10 week-old C57BL/6 mice were harvested 24h following stereotactic microinjection of murine VegfA (60 ng in 3 µL PBS) or C57BL/6 healthy mouse plasma (3 μL) or vehicle control (3µL PBS). **(K-M)** Cortical lesions were immunostained with anti-Gfap and anti-Cldn4 antibodies. **(N)** Gfap positive areas and **(O)** Cldn4 positive areas were quantified (VegfA *n*= 4, C57BL/6 healthy mouse plasma *n*=4, Vehicle control *n*=4). \**P*≤0.05, \*\**P*≤0.01, *** *P*≤0.001 Kruskal Wallis test.

To confirm this observation *in vivo*, we delivered murine VegfA or murine plasmatic proteins into the left cerebral cortex of adult mice and evaluated the consequences on Cldn4 expression by astrocytes. PBS stereotactic administration was used as a control.

*In vivo*, Gfap (Figure 5, K-N) and Cldn4 (Figure 5, K-M, O) are induced in the mice cortex, after murine VegfA and murine plasmatic protein treatments, VegfA having a much stronger effect than plasmatic proteins.

We concluded that permeable endothelial monolayers produce signals that can drive astrocyte reactivity and tight junction expression. Plasmatic protein involvement in controlling astrocyte barrier behavior is however less clear as astrocyte reactivity and tight junction expression are up regulated *in vivo* but not *in vitro* when treated with plasma and further investigations will be necessary to identify the mechanisms involved.

### Mice with endothelial Dhh inactivation display reduced disability in a model of multiple sclerosis during the onset of the disease

To examine the impact of these findings on disease severity, we investigated the phenotype of induced-experimental multiple sclerosis (EAE) in Dhh^ECKO^ and control mice.

We observed that in control mice, neurologic deficit is observed from day 9, and increases in severity until day 18, when clinical score stabilizes at a mean of 3.2, representing hind limb paralysis. In contrast, the onset of clinical signs in Dhh^ECKO^ mice is first seen 4 days later, and the clinical course is much milder. In Dhh^ECKO^ mice, disease reached a plateau at day 21 at a mean of 2.3, indicating hind limb weakness and unsteady gait, a mild phenotype (Figure 6, A). The EAE peak score (Figure 6, B) and average score during the time of disability (Figure 6, C) are both decreased in Dhh^ECKO^ mice but there are no significant changes in survival and mortality rates (Figure 6, D). Importantly, the clinical course in the Dhh^ECKO^ is correlated to strikingly decreased areas of demyelination compared to the control cohort (Figure 6, E-F). Critically, these studies revealed that the clinical course and pathology of EAE are strongly reduced in Dhh^ECKO^ mice during the onset of the disease.

**Figure 6:**
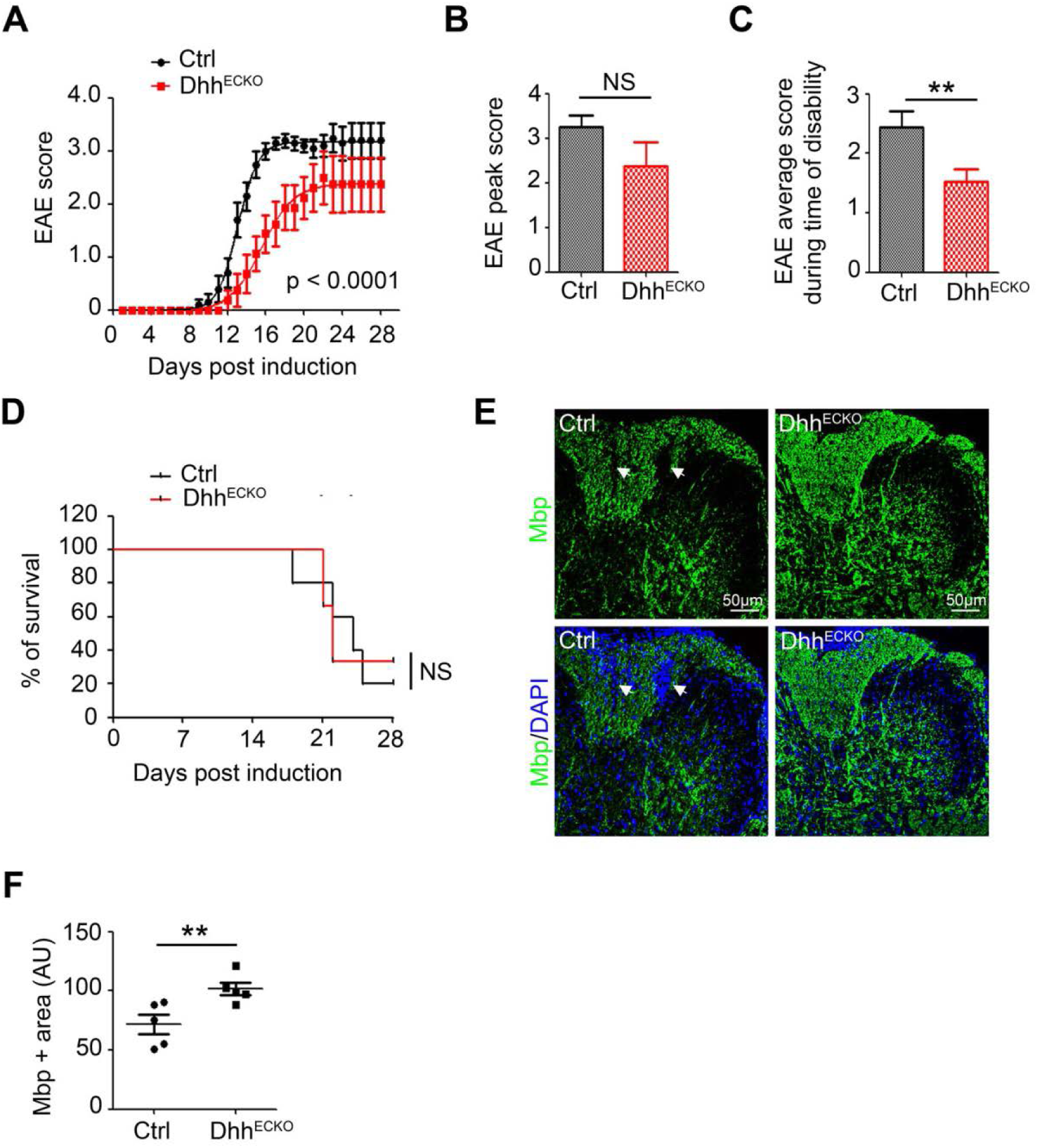
Dhh^ECKO^ mice display reduced disability in a model of multiple sclerosis during the onset of the disease: **(A)** Dhh^ECKO^ and control mice induced with EAE were scored daily on a standard 5-point scale, nonlinear regression (Boltzmann sigmoidal). **(B)** Dhh^ECKO^ and control mice EAE peak score, **(C)** EAE average score during time of disability and **(D)** mortality rate was quantified other the course of the disease. **(E-F)** Spinal cord EAE lesions from Dhh^ECKO^ mice and littermate controls were harvested at 28 days post induction or at the time of euthanasia. **(E)** Dhh^ECKO^ and control lesions were immunostained with an anti-Myelin Basic Protein (Mbp) (in green) antibody; arrows indicate white matter loss areas. Nuclei were stained with DAPI (in blue). **(F)** Mbp positive areas were quantified ((Dhh^ECKO^ *n* = 5, WT *n* = 5). \*\**P*≤0.01, Mann-Whitney U test.

We concluded that endothelial Dhh knockdown-induced BBB opening is associated to a clinical protective effect in a model of multiple sclerosis.

### Mice with endothelial Dhh inactivation display a reinforced barrier at the Glia Limitans restraining access to the parenchyma to inflammation in a model of multiple sclerosis

Although Dhh^ECKO^ mice display equivalent Fgb densities (Figure 7, A) as well as numbers of inflammatory cells (Figure 7, B) in lesions, at 28 dpi, compared to that in control mice, neuropathology in both cohorts appeared very different. We showed that while the BBB is permeable in both groups with plasmatic protein extravasation associated to equivalent Cdh5 densities, astrocyte reactivity in EAE lesions in Dhh^ECKO^ mice is greatly increased with Gfap immunoreactivity strongest at the Glia Limitans (Figure 7 C, D, E). Moreover, infiltrating plasmatic proteins and inflammatory cells in Dhh^ECKO^ mice show less CNS parenchymal dispersion (Figure 7, E-G), with 69.0 % of Fgb trapped into the Glia Limitans in the Dhh^ECKO^ cohort versus 29.8 % in the control cohort (Figure 7, E-G) and 77.1 % of Cd45+ T cells trapped into the Glia Limitans in the Dhh^ECKO^ cohort versus 25.1 % in the control cohort (Figure 7, H-J). Altogether, these data suggest less access through the Glia Limitans in the Dhh^ECKO^ mice compared to the littermate control mice. Finally we demonstrated that Cldn4 expression in spinal cord EAE lesion lysates is up regulated in Dhh^ECKO^ mice compared to control mice (Figure 7, K-L).

**Figure 7:**
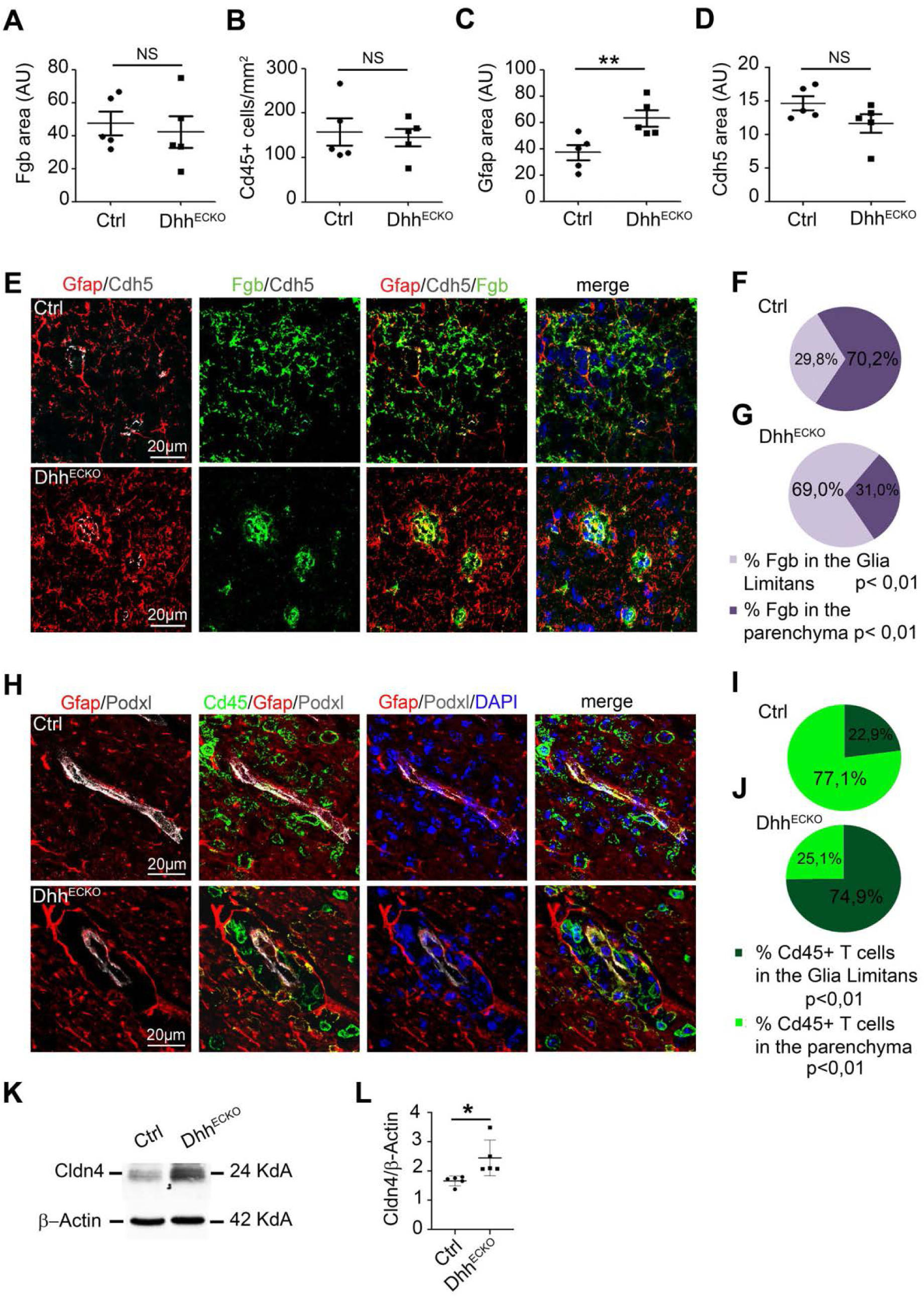
Mice with endothelial Dhh knockdown display a reinforced barrier at the Glia Limitans restraining access to the parenchyma to inflammation in a model of multiple sclerosis: **(A-I)** Spinal cord EAE lesions from Dhh^ECKO^ mice and littermate controls were harvested at 28 days post induction or at the time of euthanasia. **(A-D)** Dhh^ECKO^ and control lesions were immunostained with anti Fgb, anti-Cd45, anti-Gfap and anti-Cdh5 antibodies. **(A)** Fgb positive areas, **(B)** the number of Cd45+ T cells per mm^2^, **(C)** Gfap positive areas and **(D)** Cdh5 positive area were quantified (Dhh^ECKO^ *n* = 5, WT *n* = 5). **(E-G)** Dhh^ECKO^ and control lesions were immunostained with anti-Cdh5 (in grey), anti-Fgb (in green) and anti-Gfap (in red) antibodies (Nuclei were stained with DAPI (in blue)) and **(F-G)** the distribution of Fgb within the Glia Limitans and the parenchyma was quantified. **(H-J)** Dhh^ECKO^ and control lesions were immunostained with anti-Podxl (in grey), anti-Cd45 (in green) and anti-Gfap (in red) antibodies (Nuclei were stained with DAPI (in blue)) and **(I-J)** the distribution of Cd45+ T lymphocytes within the Glia Limitans and the parenchyma was quantified. **(K)** Cldn4 expression level was quantified by western blot on Dhh^ECKO^ and control spinal cord EAE lesion lysates. (Dhh^ECKO^ *n* = 5, WT *n* = 5). \**P*≤0.05, \*\**P*≤0.01, Mann-Whitney U test.

Collectively, data from Figure 4 to 7 reveal that conditional loss of a key structural component of endothelial integrity at the BBB in Dhh^ECKO^ mice leads to increased astrocyte reactivity and implementation of barrier properties at the Glia Limitans, allowing for less diffusion of plasmatic proteins and immune cells into the CNS parenchyma than in control mice. Therefore, in Dhh^ECKO^ mice, astrocytic endfeet at the Glia Limitans are « preconditioned” to form a barrier which explains their ability during neuropathology, to protect the parenchyma more efficiently than in the controls, leading to the protective effect observed clinically. Thus we identify BBB leakage, induced by the down regulation of Dhh endothelial expression, as an important mechanism controlling Glia Limitans reactivity and barrier properties and subsequently tissue damage and clinical deficit in a model of human disease.

## Discussion

Presently, it is accepted that the BBB is the sole line of defense of the CNS, restricting access to the parenchyma to inflammatory infiltration notably in the context of chronic neuro-inflammation. However, in our study, we brought a different perspective on CNS barrier organization, unveiling the existence of two independent, dissociable states of the astrocyte and endothelial barriers in the gliovascular unit. Indeed we confirmed that, just like the BBB, the Glia Limitans can form a protective barrier. For the first time, we demonstrated for the first time that BBB breakdown is sufficient to induce chronic barrier properties at the Glia Limitans and we highlighted a crosstalk from endothelial cells to astrocytes which restrict access to the parenchyma to plasmatic proteins and inflammatory cells during multiple sclerosis. Indeed we demonstrated that in the Dhh^ECKO^ mice which display an open BBB, there is astrocytic Cldn4 expression under resting condition, before EAE induction; therefore, with neuro-inflammation, Glia Limitans in Dhh^ECKO^ mice responds faster and more efficiently in term of barrier properties which explain that the EAE symptoms are less severe in the Dhh^ECKO^ mice compared to control littermates.

The critical role of the Hh signaling in CNS neuro-inflammation has first been highlighted in 2011 by Prat’s laboratory; this study revealed that during multiple sclerosis development, the morphogen Shh is expressed by reactive astrocytes and participates to the maintenance of BBB integrity^4^. Following this discovery, our group published in 2018 that Dhh is physiologically expressed at the BBB in adults^8^, result supported by the Betsholtz’s group study of brain mouse single cell RNA sequencing^25^. Here we demonstrated for the first time that Dhh is down regulated at the BBB during multiple sclerosis and that Dhh knockdown is sufficient to induce BBB permeability by inhibiting Cdh5 and Cldn5 expression through the modulation of FoxO1 activity, strengthening the idea that the Hh signaling is essential to control BBB integrity both physiologically and in multiple sclerosis condition. Based on our results and the literature, we may hypothesized that Dhh is necessary to maintain BBB tightness under physiological conditions and that Dhh down regulation under inflammatory conditions might be offset by astrocytic Shh secretion to maintain BBB homeostasis during disease progression.

Over the past years, the literature started to acknowledge the fact that BBB is not the sole line of defense of the CNS and that the glia that abut the vasculature might play a role in restricting access to the parenchyma. Indeed, Sofroniew’s lab first described that in spinal cord injury, astrocyte scar borders have structural similarities to the Glia Limitans borders and that functionally, astrocyte scar borders corral inflammatory cells within areas of damaged tissue^28,29^. Moreover, in 2017, we published that, during EAE multiple sclerosis development, reactive astrocytes, at the Glia Limitans, form tight junctions of their own, notably Cldn4^20^, a junction protein also expressed in tightly sealed epithelia^30,31^. Noteworthy is the fact that down regulation or reorganization of Cldns and other tight junctions has been implicated in permeability in various tissues, notably the gut^32–34^; however, reports of dynamic tight junction protein induction resulting in functional barrier formation have been rare^35,36^. Here, we highlighted for the first time that disruption of endothelial junctions is sufficient to induce Cldn4 expression at the Glia Limitans and subsequently plasmatic protein entrapment in the PVS, identifying an astrocyte inducible barrier dependent on signals transmitted by the open BBB.

It has already been described, by our group and others, that astrocytes can send signals, notably VegfA^18^, Tymp^17^ and Shh^4^, to the BBB to modulate its state (tight *versus* permeable). In this study, we acknowledge the existence of a dialogue going the opposite way, by showing that BBB endothelial junction disruption leads to Cldn4 expression at the Glia Limitans. Interestingly, endothelial cell capacity to send signals to neighboring cells has been previously identified notably in the context of pericyte (mural cells associated to arterioles, capillaries and venules) recruitment at the vascular wall^37^. Specifically, it has been shown that Pdgfb is secreted from angiogenic sprout endothelium where it serves as an attractant for co-migrating pericytes, which in turn express Pdgfrβ^38^. Based on these arguments and our results, it appears strongly consistent that endothelial signals can be sent to astrocytes. Identifying such signals will be the aim of future studies by our group.

We then showed that in animals which exhibit an open BBB (Dhh^ECKO^ mice), astrocytic endfeet at the Glia Limitans form a barrier more efficiently than in the littermate controls, leading to the protective effect observed clinically in the model of multiple sclerosis. This is somehow reminiscent of what is observed in brain ischemic preconditioning where a mild non-lethal ischemic episode (preconditioning) can produce resistance to a subsequent more severe ischemic insult^39,40^. Here, inducing BBB opening and PVS plasmatic protein accumulation produce resistance to the subsequent massive inflammatory infiltration induced by multiple sclerosis development. This may explain why multiple sclerosis relapse/lesion formation rarely occurs at the same location in the CNS. Interestingly, among neurons and non-neuronal cells, astrocytes are considered increasingly important in regulating cerebral ischemic tolerance^41^ and a parallel can be easily drawn between these results and ours showing a major role for “preconditioned” astrocytes in the control of “chronic neuro-inflammation tolerance” and protection against further relapse.

In light of the above observations, we may assume that the CNS has the ability to protect itself against isolated BBB leakage episodes through a secondary barrier at the Glia Limitans that takes over once the BBB is open. Moreover, it suggests that manipulation of the BBB and Glia Limitans in combination may have greater potential than either alone to control CNS entry of leukocytes and pro-inflammatory soluble factors, in conditions such as multiple sclerosis and perhaps more widely. Indeed, taking into account both components of the gliovascular unit is of translational interest notably to limit CNS parenchymal access to pathogenic agents by strengthening the Glia Limitans once the BBB is open, in cardiovascular diseases such as brain ischemic strokes^42^, neuro-infections^43^ and neuro-degeneration (Parkinson’s/Alzheimer’s diseases, vascular dementia)^44^ or to facilitate parenchymal access to drugs, by opening the BBB and Glia Limitans together, in CNS tumor treatment^45^. Along similar lines, it is unknown how the barrier properties of the Glia Limitans may impact the pharmacokinetics of drugs that must enter the CNS parenchyma in conditions such as multiple sclerosis, which may account for treatment failure.

In summary, our study first demonstrates the critical role of Dhh in maintaining BBB integrity. We prove that Dhh is down regulated during multiple sclerosis and that Dhh knockdown leads to BBB opening. Using Dhh knockdown as a tool to cause BBB opening, we then show that BBB permeability is sufficient to induce a secondary barrier at the Glia Limitans characterized by the expression of the tight junction Cldn4 and astrocyte reactivity. Thus highlights a double barrier system within the CNS relying on a crosstalk going from the endothelial BBB to the astrocytic Glia Limitans. Finally our study provides evidence to a novel concept of “chronic neuro-inflammation tolerance”, as stimulation of Glia Limitans barrier properties by BBB opening, upstream of multiple sclerosis development, leads to a protective effect against disease progression.

## Methods

### Human tissues

cortical sections from multiple sclerosis patients (active lesions) and healthy controls (frontal cortex) were obtained from the Neuro-CEB bio bank (https://www.neuroceb.org/fr). The sections were 30 µm thick and obtained from fresh frozen samples.

### Mice

Dhh Floxed (Dhh^Flox^) mice were generated at the “Institut Clinique de la Souris” through the International Mouse Phenotyping Consortium (IMPC) from a vector generated by the European conditional mice mutagenesis program, EUCOMM and described before^8^. Animal experiments were performed in accordance with the guidelines from Directive 2010/63/EU of the European Parliament on the protection of animals used for scientific purposes and approved by the local Animal Care and Use Committee of Bordeaux University.

The Cre recombinase in Cdh5-Cre^ERT2^ mice was activated by intraperitoneal injection of 1 mg tamoxifen (Sigma Aldrich, St. Louis, MO, USA) for 5 consecutive days at 8 weeks of age. Mice were phenotyped 2 weeks later. Successful and specific activation of the Cre recombinase has been verified before^8^. C57BL/6 mice were purchased from Jackson Laboratories (Bar Harbor, ME, USA).

### Isolation of neurovascular units from mouse CNS

Mouse was sacrificed by cervical dislocation and its head cut and rinsed with 70 % ethanol. Brain and spinal cord were then harvested and cerebellum, olfactory bulb and white matter removed from the brain with sterile forceps. Additionally, meninges were eliminated by rolling a sterile cotton swab at the surface of the cortex. The cortex and spinal cord were then transferred in a potter containing 2 mL of buffer A (HBSS 1X w/o phenol red (Gibco, Waltham, MA, USA), 10 mM HEPES (Gibco, Waltham, MA, USA) and 0,1 % BSA (Sigma Aldrich, St. Louis, MO, USA) and the CNS tissue was pounded to obtain an homogenate which was collected in a 15 mL tube. The potter was rinsed with 1 mL of buffer A which was added to the 2 mL homogenate. Cold 30 % dextran solution was then added to the tube (V:V) to obtain a 15 % dextran working solution centrifuged for 25 minutes at 3000 g, 4 °C without brakes. After centrifugation, the pellet (neurovascular components and red cells) was collected and the supernatant (dextran solution and neural components) was centrifuged again to get the residual vessels. Neurovascular components were then pooled and re-suspended in 4 mL of buffer B (HBSS 1X Ca^2+^/ Mg^2+^ free with phenol red (Gibco, Waltham, MA, USA), 10 mM HEPES (Gibco, Waltham, MA, USA) and 0,1 % BSA (Sigma Aldrich, St. Louis, MO, USA)).

#### Isolation of neurovascular units for RT-PCR, western blots or immunohistochemistry

after centrifugation of the cell suspension, the pellet was washed 3 times with the buffer B and filtered through a 100 µm nylon mesh (Millipore Corporation, Burlington, MA, USA). The nylon mesh was washed with 7 mL of buffer B to collect the retained neurovascular units. The suspension was then centrifuged for 10 minutes at 1000 g and the pellet suspended in 300 µL of RIPA lysis buffer for western blot analysis or 1000 µL of Tri-Reagent (MRC, Cincinnati, OH, USA) for RTqPCR analysis. For immunohistochemistry, the pellet was suspended in 3 mL of a solution of matrigel (Corning, Steuben, NY, USA)-DMEM 1 g/L glucose, Mg^+^, Ca^2+^ (Gibco, Waltham, MA, USA) 1:80, distributed on a labtek (Starstedt, Nümbrecht, Germany) (one mouse brain is needed to seed 1 labtek) and incubated for 30 minutes at 37 °C. Finally the neurovascular units embedded in the matrigel (Corning, Steuben, NY, USA) solution were fixed with 10 % formalin for 10 minutes.

#### Primary culture of mouse CNS micro vascular endothelial cells (CNS MECs)

after centrifugation of the cell suspension, the pellet was washed 3 times with the buffer B and transferred in an enzyme solution (2 mg/mL Collagenase/Dispase (Roche, Bale, Switzerland), 0,147 µg/mL TLCK (Lonza, Bäle, Switzerland) and 10 µg/mL DNAse 1 (Roche, Bäle, Switzerland)), pre-warmed at 37 °C, before being placed on a shaking table at maximum speed agitation at 37 °C. After 30 minutes, the digestion was stopped by adding 10 mL of buffer B, the cell suspension centrifuged and the digested neurovascular pellet washed 3 times with 3 mL of buffer B. After the 3 washing steps, the digested neurovascular pellet was re-suspended in 1 mL of Mouse Brain Endothelial Cell Culture Medium (DMEM 1 g/L glucose, Mg^+^, Ca^2+^ (Gibco, Waltham, MA, USA), FBS 20 % (Gibco, Waltham, MA, USA), sodium pyruvate 2 % (Gibco, Waltham, MA, USA), non-essential amino acids 2 % (Lonza, Bäle, Switzerland), FGF 1 ng/mL (PeproTech, Rocky hill, NJ, USA) and gentamycin 10 mg/mL (Gibco, Waltham, MA, USA)) and plated on a labtek (Starstedt, Nümbrecht, Germany) (1 mouse brain is needed to seed 1 labtek) previously coated with 2 % matrigel (Corning, Steuben, NY, USA) diluted in DMEM 1 g/L glucose, Mg^+^, Ca^2+^ (Gibco, Waltham, MA, USA). **Cell culture:** Human Brain Microvascular Endothelial Cells (HBMECs) (Alphabioregen-CliniSciences, Nanterre, France) were cultured in endothelial basal medium-2 (EBM-2) supplemented with EGM™-2 BulletKits™ (Lonza, Bäle, Switzerland). Cell from passage 3 to passage 6 were used. Before any treatment, cells were serum starved in 0.5 % fetal bovine serum EGM-2 medium for 24 hours. Normal Human Astrocytes (NHA) (Lonza, Bäle, Switzerland) were cultured in astrocyte basal medium (ABM) supplemented with AGM™ BulletKits™ (Lonza, Bäle, Switzerland). Cell from passage 2 to passage 4 were used. Before any treatment, cells were serum starved in DMEM 1 g/L glucose, Mg^+^, Ca^2+^ (Gibco, Waltham, MA, USA) without serum for 24 hours.

### Cytokines/Growth Factors/Chemicals

Human IL-1β was purchased from PeproTech (Rocky Hills, NJ, USA) and Human and mouse Vegf-165 (VegfA) were purchased from CliniSciences (Nanterre, France). Based on previous studies, Human IL-1β and Human Vegf-165 were routinely used at 10 ng/mL^19,21^. Mouse Vegf-165 was used at a concentration of 20 ng/µL. The inhibitor of FoxO1 total (AS1842856) was purchased from Merck (Kenilworth, NJ, USA) and was used at 100 nM^22^. D-Mannitol was purchased from Sigma Aldrich (Saint Louis, MI, USA) and was used at 100 mM^23^.

### Antibodies

Anti-Cldn4 (mouse), anti-Cldn5 (mouse (tissues) and rabbit (cell culture)), anti-human Zo1 (rabbit) and anti-Gfap (rat) were from Invitrogen (Carlsbad, CA, USA). Anti-Cdh5 (goat) and anti-Podxl (goat) were from R&D systems (Minneapolis, MN, USA). Anti-human Cdh5 (mouse) and anti-human Dhh (H-85) (rabbit) were from Santa Cruz Biotech (Santa Cruz, CA, USA). Anti-Fgb (rabbit) and anti-human Pecam1 (mouse) were from Dako (Carpinteria, CA, USA). Anti-Alb (sheep) and anti-Mbp (rat) were from Abcam (Cambridge, MA, USA). Anti-Cd4, Cd11b, and Cd45 (rat) were from eBioscience (San Diego, CA, USA). Anti-NeuN (rabbit) was from Millipore (Billerica, MA, USA). Anti-Lam (rabbit) was from Sigma Aldrich (St. Louis, MO, USA). Anti-Zo1 (rabbit) was from Life Technologies (Grand Island, NY, USA). Anti-FoxO1 (rabbit) anti-p-FoxO1 (rabbit) and anti-β-actin (rabbit) were from cell signaling (Danvers, MA, USA).

### Quantitative RT-PCR

RNA was isolated using Tri Reagent® (Molecular Research Center Inc) as instructed by the manufacturer, from 3×10^5^ cells or from isolated mouse CNS neurovascular units. For quantitative RT-PCR analyses, total RNA was reverse transcribed with M-MLV reverse transcriptase (Promega, Madison, WI, USA) and amplification was performed on a DNA Engine Opticon®2 (MJ Research Inc, St Bruno, Canada) using B-R SYBER® Green SuperMix (Quanta Biosciences, Beverly, MA, USA). Primer sequences are reported in Supplementary table 1.

**Table 1:**
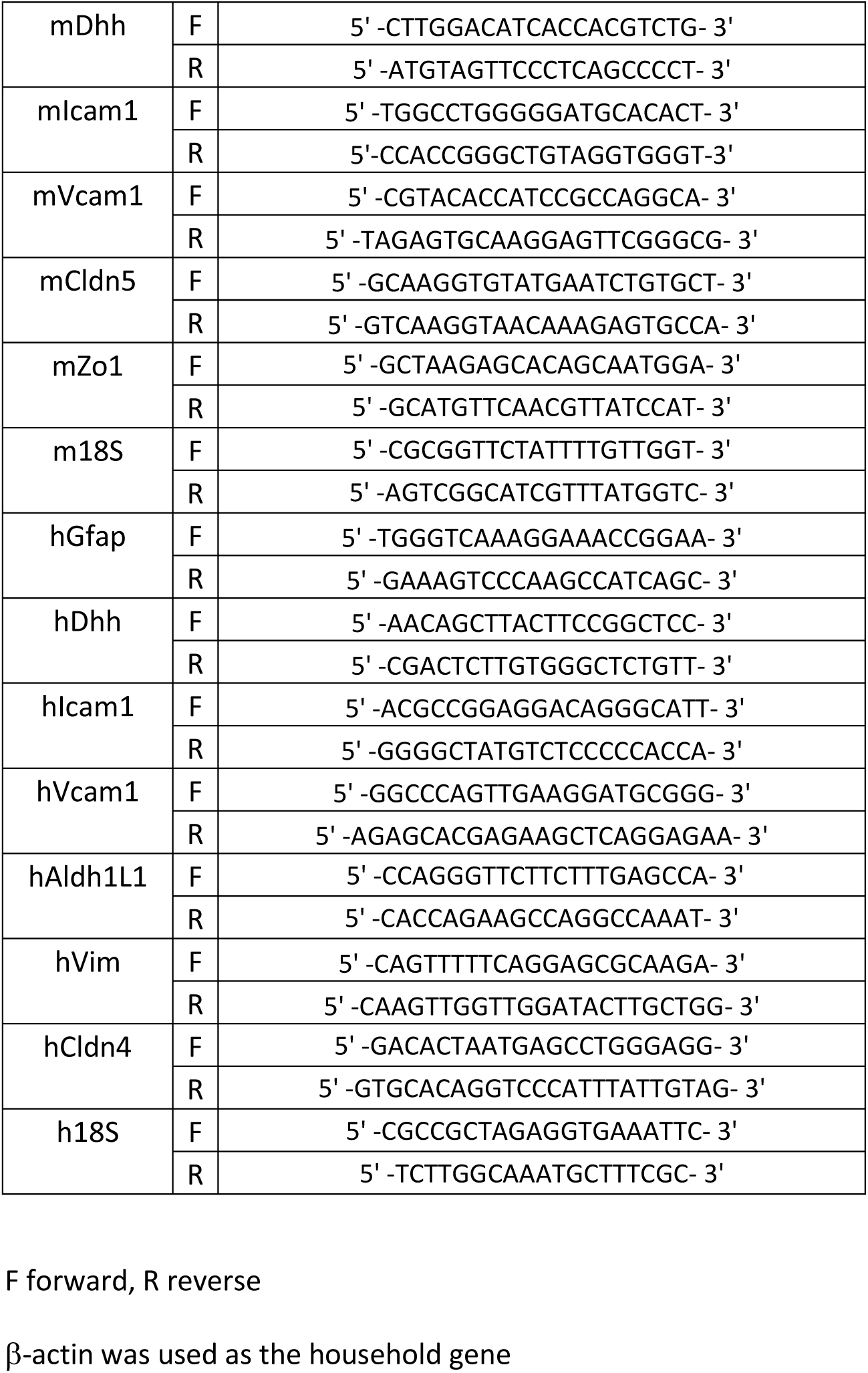
List of primers used for reverse transcription (RT) quantitative polymer chain reaction (qPCR)

The relative expression of each mRNA was calculated by the comparative threshold cycle method and normalized to HPRT mRNA expression.

### Western Blots

Protein expression was evaluated by SDS-PAGE. Protein loading quantity was controlled using the rabbit monoclonal anti-β-actin antibody (cell signaling, Danvers, MA, USA). Secondary antibodies were from Invitrogen. The signal was then revealed by using an Odyssey Infrared imager (LI-COR, Lincoln, NE, USA). For quantification, the mean pixel density of each band was measured using Image J software (NIH, Bethesda, MD, USA) and data were standardized to β-actin, and fold change versus control calculated.

### Stereotactic injection

10 weeks old C57BL/6 mice (4 mice per condition) were anesthetized and secured into a stereotactic frame (Stoelting Co., IL, USA). 3μl of murine VegfA (20 ng/µL), 3 µL of healthy mouse plasma or 3 µL of vehicle control (PBS) were delivered at 0.1 μl/s into the frontal cortex at coordinates of 1 um posterior to bregma, 2 µm left of the midline and 1.5 um below the surface of the cortex. Mice were sacrificed by pentobarbital (Richter Pharma, Wels, Austria) overdose at 24h post injection (dpi). For histological assessment, the brain of each animal was harvested.

### Experimental autoimmune encephalomyelitis (EAE)

10 week old female mice were immunized by subcutaneous injection of 300 µg myelin oligodendrocyte glycoprotein-35-55 (MOG_35–55_)(Hooke laboratories, Lawrence, MA, USA) in 200 µl Freund’s Adjuvant containing 300 µg/mL mycobacterium tuberculosis H37Ra (Hooke laboratories, Lawrence, MA, USA) in the dorsum. Mice were administered with 500 ng pertussis toxin (PTX) intra*-*peritoneously on day of sensitization and 1 day later (Hooke laboratories). The emulsion provides antigen which initiates expansion and differentiation of MOG-specific autoimmune T cells. PTX enhances EAE development by providing additional adjuvant. EAE will develop in mice 7-14 days after immunization (Day 0): animals which develop EAE will become paralyzed. Disease was scored (0, no symptoms; 1, floppy tail; 2, hind limb weakness (paraparesis); 3, hind limb paralysis (paraplegia); 4, fore- and hind limb paralysis; 5, death)^24^ from day 7 post immunization until day 32 post immunization. At Day 32, all the animals were euthanized by pentobarbital (Richter Pharma, Wels, Austria) overdose. For histological assessment, cervical, lumbar and dorsal sections of each animal spinal cord, as well as the spleen, were harvested.

### Immunohistochemistry

prior to staining, brain and spinal cord samples were either fixed in 10 % formalin for 3 hours, incubated in 30 % sucrose overnight, OCT embedded and cut into 9 µm thick sections or directly OCT embedded and cut into 9 µm thick sections. Cultured cells were fixed with 10 % formalin for 10 minutes. Human frozen sections were used directly without any prior treatment. Concerning the fixed sections, for Cldn4, prior to blocking, sections were soaked in Citrate (pH 7.5; 100 °C). For Cldn5, prior to blocking, sections were soaked in EDTA (pH 6.0; 100 °C). For Cd4, Cd11b, and Cd45, sections were treated with 0.5 mg/mL protease XIV (Sigma Aldrich, St. Louis, MO, USA) at 37 °C for 5 minutes. Primary antibodies were used at 1:100 except Cldn4 (1:50), Fgb (1:1,000), and Alb (1:1,000). Samples were examined using a Zeiss Microsystems confocal microscope (Oberkochen, Germany), and stacks were collected with *z* of 1 μm.

For immunofluorescence analyzes, primary antibodies were resolved with Alexa Fluor®–conjugated secondary polyclonal antibodies (Invitrogen, Carlsbad, CA, USA) and nuclei were counterstained with DAPI (1:5000) (Invitrogen, Carlsbad, CA, USA). For all immunofluorescence analyses, negative controls using secondary antibodies only were done to check for antibody specificity.

### Morphometric analysis

Morphometric analyses were carried out using NIH ImageJ software (NIH, Bethesda, MD, USA).

*BBB permeability* was evaluated by measuring tight junction integrity and plasmatic protein extravasation. Brain and spinal cord sections were immunostained for the expression of Cldn5/Cdh5 and Fgb/IgG/Alb respectively. For each brain or spinal cord section, Cldn5+, Cdh5+, Fgb+, IgG+ and Alb+ areas were quantified in 20 pictures taken at the margins of the lesion area under 40x magnification. One section was quantified per spinal cord (3 different zones are displayed on the same section: one cervical, one lumbar and one dorsal to get a global vision of the lesion) (per mouse).

*Leukocyte, lymphocyte and macrophage densities* were evaluated in sections stained for the expression of Cd45, Cd4 and Cd11b respectively. For each brain or spinal cord section, Cd45+ leukocytes, Cd4+ lymphocytes and CD11b+ macrophages were counted in 20 pictures randomly taken under 40x magnification. One section was quantified per spinal cord (3 different zones are displayed on the same section: one cervical, one lumbar and one dorsal to get a global vision of the inflammatory lesion) (per mouse).

*Demyelination* was evaluated in spinal cord sections stained for the expression of Mbp. For each spinal cord section, Mbp+ area was quantified in 10 pictures taken in and around inflammatory lesion sites under 20X magnification. One section was quantified per spinal cord (3 different zones are displayed on the same section: one cervical, one lumbar and one dorsal to get a global vision of the lesion) (per mouse).

*Plasmatic protein and leukocyte infiltrate distribution at the gliovascular unit* were evaluated in spinal cord sections triple-stained for Cdh5 or Podxl (markers of the BBB), Fgb, IgG (plasmatic proteins) or Cd45 (leukocyte infiltrate) and Gfap (marker of astrocyte reactivity). For each spinal cord section, the distribution (between the lumen, the PVS and the parenchyma) of the Fgb/IgG or the leukocyte infiltrate was quantified for 2 gliovascular units in 5 pictures randomly taken under 40x magnifications. One section was quantified per spinal cord (3 different zones are displayed on the same section: one cervical, one lumbar and one dorsal to get a global vision of the lesion) (per mouse).

### Statistical analyses

Results are reported as mean ± SEM. Comparisons between groups were analyzed for significance with the non-parametric Mann-Whitney test, the non parametric Kruskal Wallis test followed by the Dunn’s multiple comparison test when we have more than 2 groups or a nonlinear regression test (Boltzmann sigmoidal) for the EAE scoring analysis using GraphPad Prism v8.0.2 (GraphPad Inc, San Diego, CA, USA). Differences between groups were considered significant when p≤0.05 (*: p≤0.05; **: p≤0.01; ***: p≤0.001).

## Author contributions

P.-L. H. and S. G. conducted experiments, acquired data, analyzed data P.M., A. D. and L.C. conducted experiments, acquired data. A.-P. G., T. C. and M.-A. R. critically revised the manuscript. C.C. designed research studies, conducted experiments, acquired data, analyzed data, provided reagents, and wrote the manuscript

## Acknowledgments

We thank Annabel Reynaud, Sylvain Grolleau, and Maxime David for their technical help. We thank Christelle Boullé for administrative assistance.

This study was supported by grants from the European Concil (Marie Skłodowska-Curie Actions, Individual fellowship 2019 (MSCA-IF-2019)) and the Fondation ARSEP (Fondation pour la Recherche sur la Sclérose En Plaques). This study was also co-funded by the “Institut National de la Santé et de la Recherche Médicale” and by the University of Bordeaux.

## Abbreviations

Alb: Albumin
Aldh1l1: Aldehyde dehydrogenase 1 family, member L1
BBB: Blood Brain Barrier
Cd4: Cluster of differentiation 4
Cd45: Cluster of differentiation 45
Cdh5: Cadherin5
Cldn4: Claudin4
Cldn5: Claudin5
Dhh: Desert hedgehog
FoxO1: Forkhead box O1
Fgb: Fibrinogen beta chain
Gfap: Glial fibrillary acidic protein
Icam1: Intercellular adhesion molecule 1
IgG: Immunoglobulin G
IL-1β: Interleukin-1β
Mbp: Myelin basic protein
NeuN: b Platelet derived growth factor subunit b
Pdgfrβ: Platelet derived growth factor receptor beta
Pecam1: Platelet/endothelial cell adhesion molecule 1
Podxl: Podocalyxin
PVS: Perivascular space
Tymp: Thymidine phosphorylase
Vcam1: Vascular cell adhesion molecule 1
VegfA: Vascular endothelial growth factor A
Vim: Vimentin
Zo1: Zonula Occludens 1

## Bibliography

1. McConnell, H. L., Kersch, C. N., Woltjer, R. L. & Neuwelt, E. A. The Translational Significance of the Neurovascular Unit. J. Biol. Chem. 292, 762–770 (2017).

2. Bartholomäus, I. et al. Effector T cell interactions with meningeal vascular structures in nascent autoimmune CNS lesions. Nature 462, 94–98 (2009).

3. Brück, W. et al. Inflammatory central nervous system demyelination: correlation of magnetic resonance imaging findings with lesion pathology. Ann. Neurol. 42, 783–793 (1997).

4. Alvarez, J. I. et al. The Hedgehog pathway promotes blood-brain barrier integrity and CNS immune quiescence. Science 334, 1727–1731 (2011).

5. Singh, V. B., Singh, M. V., Gorantla, S., Poluektova, L. Y. & Maggirwar, S. B. Smoothened Agonist Reduces Human Immunodeficiency Virus Type-1-Induced Blood-Brain Barrier Breakdown in Humanized Mice. Sci. Rep. 6, 26876 (2016).

6. Xia, Y. et al. Recombinant human sonic hedgehog protein regulates the expression of ZO-1 and occludin by activating angiopoietin-1 in stroke damage. PloS One 8, e68891 (2013).

7. Chapouly, C., Guimbal, S., Hollier, P.-L. & Renault, M.-A. Role of Hedgehog Signaling in Vasculature Development, Differentiation, and Maintenance. Int. J. Mol. Sci. 20, (2019).

8. Caradu, C. et al. Restoring Endothelial Function by Targeting Desert Hedgehog Downstream of Klf2 Improves Critical Limb Ischemia in Adults. Circ. Res. 123, 1053–1065 (2018).

9. Bitgood, M. J. & McMahon, A. P. Hedgehog and Bmp genes are coexpressed at many diverse sites of cell-cell interaction in the mouse embryo. Dev. Biol. 172, 126–138 (1995).

10. Robbins, D. J., Fei, D. L. & Riobo, N. A. The Hedgehog signal transduction network. Sci. Signal. 5, re6 (2012).

11. Iadecola, C. The Neurovascular Unit Coming of Age: A Journey through Neurovascular Coupling in Health and Disease. Neuron 96, 17–42 (2017).

12. Sweeney, M. D., Ayyadurai, S. & Zlokovic, B. V. Pericytes of the neurovascular unit: key functions and signaling pathways. Nat. Neurosci. 19, 771–783 (2016).

13. Cheslow, L. & Alvarez, J. I. Glial-endothelial crosstalk regulates blood-brain barrier function. Curr. Opin. Pharmacol. 26, 39–46 (2016).

14. Engelhardt, B. & Coisne, C. Fluids and barriers of the CNS establish immune privilege by confining immune surveillance to a two-walled castle moat surrounding the CNS castle. Fluids Barriers CNS 8, 4 (2011).

15. Engelhardt, B. & Ransohoff, R. M. Capture, crawl, cross: the T cell code to breach the blood-brain barriers. Trends Immunol. 33, 579–589 (2012).

16. Colombo, E. & Farina, C. Astrocytes: Key Regulators of Neuroinflammation. Trends Immunol. 37, 608–620 (2016).

17. Chapouly, C. et al. Astrocytic TYMP and VEGFA drive blood-brain barrier opening in inflammatory central nervous system lesions. Brain J. Neurol. 138, 1548–1567 (2015).

18. Argaw, A. T. et al. Astrocyte-derived VEGF-A drives blood-brain barrier disruption in CNS inflammatory disease. J. Clin. Invest. 122, 2454–2468 (2012).

19. Argaw, A. T., Gurfein, B. T., Zhang, Y., Zameer, A. & John, G. R. VEGF-mediated disruption of endothelial CLN-5 promotes blood-brain barrier breakdown. Proc. Natl. Acad. Sci. U. S. A. 106, 1977–1982 (2009).

20. Horng, S. et al. Astrocytic tight junctions control inflammatory CNS lesion pathogenesis. J. Clin. Invest. 127, 3136–3151 (2017).

21. Liu, J., Zhao, M. L., Brosnan, C. F. & Lee, S. C. Expression of type II nitric oxide synthase in primary human astrocytes and microglia: role of IL-1beta and IL-1 receptor antagonist. J. Immunol. Baltim. Md 1950 157, 3569–3576 (1996).

22. Karki, S. et al. Forkhead box O-1 modulation improves endothelial insulin resistance in human obesity. Arterioscler. Thromb. Vasc. Biol. 35, 1498–1506 (2015).

23. Hess, D. C. et al. Hypertonic mannitol loading of NF-kappaB transcription factor decoys in human brain microvascular endothelial cells blocks upregulation of ICAM-1. Stroke 31, 1179–1186 (2000).

24. Gurfein, B. T. et al. IL-11 regulates autoimmune demyelination. J. Immunol. Baltim. Md 1950 183, 4229–4240 (2009).

25. He, L. et al. Single-cell RNA sequencing of mouse brain and lung vascular and vessel-associated cell types. Sci. Data 5, 180160 (2018).

26. Taddei, A. et al. Endothelial adherens junctions control tight junctions by VE-cadherin-mediated upregulation of claudin-5. Nat. Cell Biol. 10, 923–934 (2008).

27. Davalos, D. & Akassoglou, K. Fibrinogen as a key regulator of inflammation in disease. Semin. Immunopathol. 34, 43–62 (2012).

28. Wanner, I. B. et al. Glial scar borders are formed by newly proliferated, elongated astrocytes that interact to corral inflammatory and fibrotic cells via STAT3-dependent mechanisms after spinal cord injury. J. Neurosci. Off. J. Soc. Neurosci. 33, 12870–12886 (2013).

29. Sofroniew, M. V. Astrocyte barriers to neurotoxic inflammation. Nat. Rev. Neurosci. 16, 249–263 (2015).

30. Hewitt, K. J., Agarwal, R. & Morin, P. J. The claudin gene family: expression in normal and neoplastic tissues. BMC Cancer 6, 186 (2006).

31. Acharya, P. et al. Distribution of the tight junction proteins ZO-1, occludin, and claudin−4, −8, and −12 in bladder epithelium. Am. J. Physiol. Renal Physiol. 287, F305–318 (2004).

32. Argaw, A. T., Gurfein, B. T., Zhang, Y., Zameer, A. & John, G. R. VEGF-mediated disruption of endothelial CLN-5 promotes blood-brain barrier breakdown. Proc. Natl. Acad. Sci. U. S. A. 106, 1977–1982 (2009).

33. Bruewer, M. et al. Proinflammatory Cytokines Disrupt Epithelial Barrier Function by Apoptosis-Independent Mechanisms. J. Immunol. 171, 6164–6172 (2003).

34. Kebir, H. et al. Human TH17 lymphocytes promote blood-brain barrier disruption and central nervous system inflammation. Nat. Med. 13, 1173–1175 (2007).

35. Kinugasa, T., Sakaguchi, T., Gu, X. & Reinecker, H. Claudins regulate the intestinal barrier in response to immune mediators. Gastroenterology 118, 1001–1011 (2000).

36. Fujita, H. et al. Claudin-1 expression in airway smooth muscle exacerbates airway remodeling in asthmatic subjects. J. Allergy Clin. Immunol. 127, 1612-1621.e8 (2011).

37. Gaengel Konstantin, Genové Guillem, Armulik Annika & Betsholtz Christer. Endothelial-Mural Cell Signaling in Vascular Development and Angiogenesis. Arterioscler. Thromb. Vasc. Biol. 29, 630–638 (2009).

38. Lindahl, P., Johansson, B. R., Levéen, P. & Betsholtz, C. Pericyte loss and microaneurysm formation in PDGF-B-deficient mice. Science 277, 242–245 (1997).

39. Yunoki, M. et al. Ischemic Tolerance of the Brain and Spinal Cord: A Review. Neurol. Med. Chir. (Tokyo) 57, 590–600 (2017).

40. Kirino, T., Tsujita, Y. & Tamura, A. Induced tolerance to ischemia in gerbil hippocampal neurons. J. Cereb. Blood Flow Metab. Off. J. Int. Soc. Cereb. Blood Flow Metab. 11, 299–307 (1991).

41. Koizumi, S., Hirayama, Y. & Morizawa, Y. M. New roles of reactive astrocytes in the brain; an organizer of cerebral ischemia. Neurochem. Int. 119, 107–114 (2018).

42. Cai, W. et al. Dysfunction of the neurovascular unit in ischemic stroke and neurodegenerative diseases: An aging effect. Ageing Res. Rev. 34, 77–87 (2017).

43. Hou, J., Baker, L. A., Zhou, L. & Klein, R. S. Viral interactions with the blood-brain barrier: old dog, new tricks. Tissue Barriers 4, e1142492 (2016).

44. Zlokovic, B. V. The blood-brain barrier in health and chronic neurodegenerative disorders. Neuron 57, 178–201 (2008).

45. Chacko, A.-M. et al. Targeted delivery of antibody-based therapeutic and imaging agents to CNS tumors: crossing the blood-brain barrier divide. Expert Opin. Drug Deliv. 10, 907–926 (2013).

